# Effect of sinusoidal electrical cortical stimulation on brain cells

**DOI:** 10.1101/855395

**Authors:** Seungjun Ryu, Kyung-Tai Kim, Hyeon Seo, Jongwook Cho, Jiyoung Park, Sung Chan Jun, Hyoung-Ihl Kim

## Abstract

**Background:** Electrical cortical stimulation is often used in patients with neurological disorders but it is unclear how it modulates different types of brain cells.

**Objective:** The aim of this study was to determine the effect of sinusoidal electrical brain stimulation (SEBS) on different types of brain cells and to identify the exact types of brain cells that are stimulated.

**Methods:** The study subjects were 40 male Sprague Dawley rats (weight 300–350 g; age 9 weeks). SEBS was delivered continuously at frequencies of 20, 40, 60, or 100 Hz to the sensory parietal cortex using epidurally placed electrodes for 1 week. Transverse rat brain tissue sections were immunolabeled with calmodulin-dependent protein kinase II and parvalbumin (PV) antibodies and with c-Fos for counting of activated excitatory and inhibitory neurons. Computer simulation was performed to cross-validate the frequency-specific cell stimulation results.

**Results:** Inhibitory neurons were more excited than excitatory neurons after epidural EBS. Most excitatory neural activity was evoked at 40 Hz (*p*<0.05) and most inhibitory neuronal activity was evoked at 20 Hz (*p*<0.01). The contralateral sensory cortex was activated significantly more at 40 Hz (*p*<0.05) and the corticothalamic circuit at 20 Hz (*p*<0.001). Stimulation-induced excitatory and inhibitory neuronal activation was widest at 20 Hz.

**Conclusions:** Epidural electrical stimulation targets both excitatory and inhibitory neurons and the related neural circuits. Further exploration is needed to identify circuits that promote the plasticity needed for recovery in patients with specific neurological diseases.

**Graphical Abstract:** 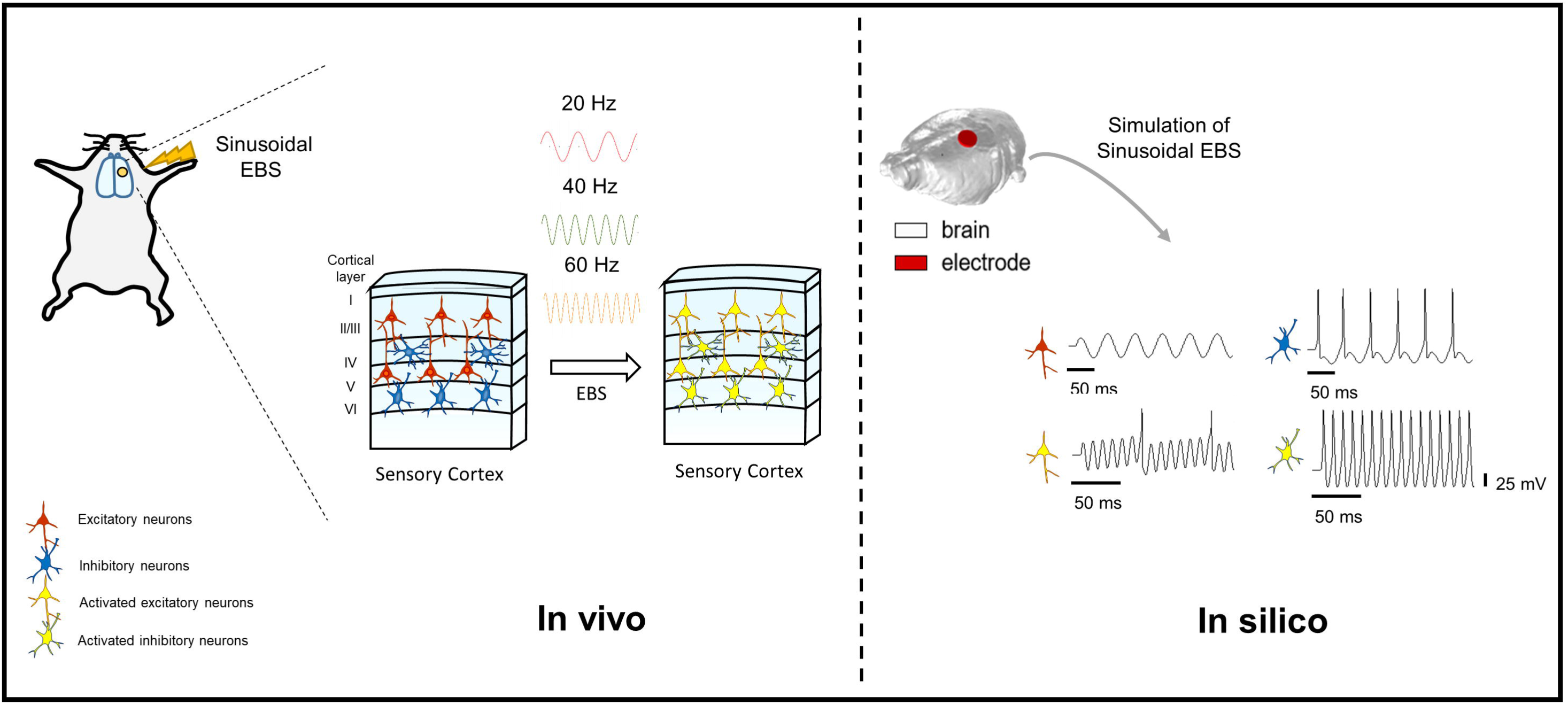

## INTRODUCTION

Electrical brain stimulation (EBS) is a type of electrotherapy that modulates neuronal activity using a controlled electric current and is increasingly used alone or in combination with other clinical therapy for various neurological disorders, such as essential tremor [1, 2], epilepsy [3, 4], Parkinson’s disease [5, 6], chronic pain [7], depression [8], cerebral infarction, and other brain disorders [9, 10]. Essentially, EBS generates an electrical field that affects specific populations of neurons. Numerous researchers have attempted to determine how this electrical field influences neural activity in local and remote neural circuits and the casual relationship with the resulting behavioral changes. However, there are many different types of neuronal cells that are under the influence of specific neurotransmitters, and it is still unclear exactly how EBS works.

The specific function triggered when a neural circuit is stimulated depends on the type of brain cell involved. Each type of neuron has a specific modulatory effect, such as post-synaptic excitation or inhibition, and the different types of neuron are interconnected. Several studies have shown that the efficacy of transmission at the synapse can undergo a short-term increase (known as facilitation) or decrease (depression) according to the activity of the presynaptic neuron [11–14]. However, previous studies of the mechanism of EBS have usually focused on the function of pyramidal neurons [15–18], given that they are assumed to play a key role in activation of neural activity and the associated plasticity in the stimulated cortex. Although recent studies have shown that excitatory neurons are strongly regulated by inhibitory neurons via feed-forward and feedback mechanisms [19, 20], the type of cells most influenced by EBS is still unknown [21–23].

The basic mechanisms underlying EBS include functional reorganization of neural structures, substrates, and increased synaptic plasticity, which are modified by various factors, such as the stimulation type and parameter, and current brain status [24–26]. Recently, transcranial alternating current stimulation(tACS) has become popular because it shows entrainment of brain oscillations in a frequency-specific manner and can be administered using various parameters, including sinusoidal weak intensity stimulation [23], but there is doubt regarding their effectiveness [27]. And also, there remain many uncertainties regarding the interaction between neural excitability and strong sinusoidal stimulation.

The aim of this study was to identify the neural cell populations that are activated during SEBS. Computational models were incorporated to clarify the effect of SEBS on the relationship between the spatial distribution of a stimulus-induced electrical field and activation of individual neurons and how it alters neuronal spiking.

## MATERIAL AND METHODS

### Experimental animals

Forty male Sprague Dawley rats (300–350 g, aged 9 weeks) were used in the study. All experiments were performed in accordance with the ARRIVE guidelines and the institutional guidelines of the Gwangju Institute of Science and Technology (GIST). All procedures were approved by the Institutional Animal Care and Use Committee at GIST. The rats were divided into four experimental groups (to receive SEBS at 20, 40, 60, or 100 Hz) and a sham operation group. At least 5 rats were included in each study group.

### Surgical procedures

The rats were anesthetized using a mixture of ketamine hydrochloride 100 mg/kg and xylazine 7 mg/kg. After 15 minutes, the rats were fixed in a small-animal stereotactic frame. Body temperature was maintained at 37.5±5°C with a thermocouple blanket. With bregma (B) and lambda (L) in a flat plane as reference points, a small craniectomy was performed 3 mm posterior to bregma and 3 mm lateral to the midline. All 40 rats underwent insertion of a custom-made electrode (diameter 3 mm, height 0.37 mm) via craniotomy in the epidural area, with a 0.7-mm-diameter reference screw electrode placed 2 mm anterior to bregma and 3 mm lateral to the midline. The electrode extended from 1.5 mm to 4.5 mm posteriorly and 1.5 mm to 4.5 mm lateral to bregma (9 mm^2^), covering the hindlimb, trunk, and forelimb areas of the sensory cortex. The electrodes were connected to a pedestal on the skull, fixed and sealed with bone cement, and then connected to a stimulator (Cybermedic Co. Ltd., Iksan, Korea) via a swivel adaptor at the top of the cage.

### Electrical stimulation

Voltage stimulation was delivered continuously (24 h/day) to the sensory cortex via a programmable Cybermedic stimulator for 1 week. We maintained the experimental stimulation intensity at half of the individual movement threshold. On alternate days, we measured the individual motor threshold during stimulation and regulated the voltage. The experimental stimulation intensities ranged from 1.0 V to 3.0 V and frequencies of 20, 40, 60, and 100 Hz were used to investigate differential stimulation of neuronal cells. A continuous sinusoidal waveform with a duty cycle of 99% was maintained for all animals in each of the experimental groups (Fig. 1A).

**Figure 1.**
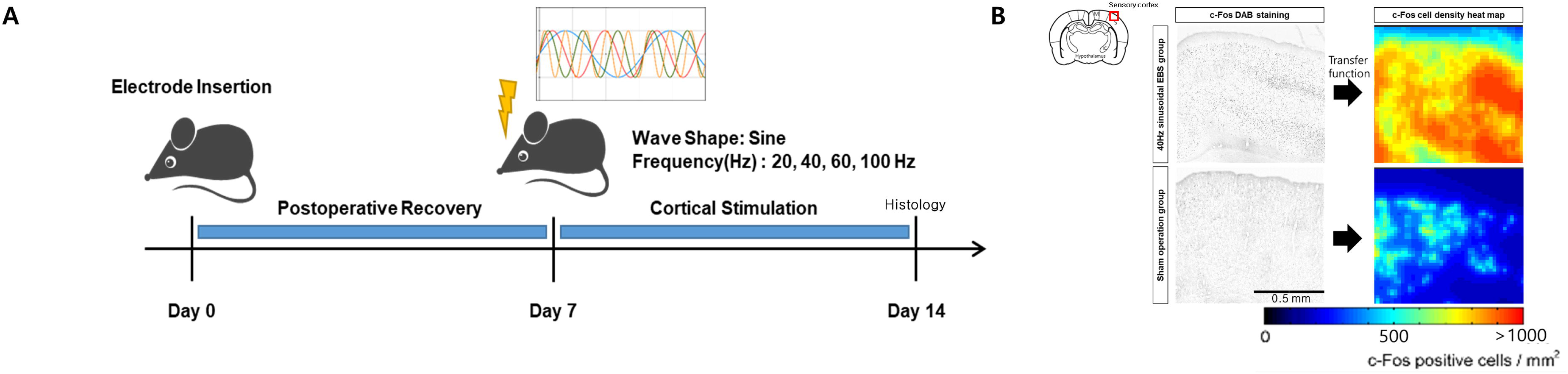
Schematic diagram of the experimental protocol and an example of cell density heat map of c-Fos-positive cells. (A) Experimental procedure for SEBS in Sprague Dawley rats. (B) Image showing the customized transfer function for cell density maps of c-Fos-positive cells. EBS, electrical brain stimulation

### Neurohistological analysis

#### Immunohistochemistry

All rats in each group were euthanized and processed for c-Fos immunohistochemistry after 1 week of cortical stimulation. Immunohistochemistry was performed as described previously [28]. Briefly, rats were perfused with 4% paraformaldehyde (PFA), post-fixed in 4% PFA overnight, and the brains were sunk to 30% sucrose for cryoprotection. Coronal brain sections (40µm) were performed using microtome. Rat brain sections were incubated with following antibodies: rabbit anti-c-Fos (1:1000) (Cell Signaling, 2250S), Mouse anti-CamKII (Abcam, ab22609), Guinea pig anti-parvalbumin (Synaptic systems, 195 004), goat anti-guinea pig alexa 555 (1:200) (Invitrogen, A21435). Proper fluorophore-conjugated secondary antibody (Invitrogen) was used and images were captured using LSM-800 confocal microscope (Zeiss).

For DAB staining, the brain sections were treated with 3% H_2_O_2_ in Tris-buffered saline and 1% normal goat serum and then incubated in c-Fos 9F6 rabbit antibody (Cell Signaling Technology). The sections were incubated in a Polink-1 horseradish peroxidase detection system for rabbit antibody (GBI Labs, Mukilteo, WA, USA) on the following day. After a color reaction was observed on incubating sections with diaminobenzidine/peroxidase solution (DAB 0.02%; 0.08% nickel sulfate) in Tris-buffered saline, the brain sections were mounted on gelatin-coated slides. c-Fos images were captured using a Leica microscope.

#### c-Fos mapping

Fast Fourier transform-bandpass filtered images were created in ImageJ (National Institutes of Health, Bethesda, MD, USA) and cell density maps using a custom MATLAB-based program (MathWorks, Natick, MA, USA) [29]. A sample image showing the results of the transfer function when applied is shown in Fig. 1B. Regions of interest were selected on the motor and sensory cortices, striatum, and thalamus, and the number of c-Fos-positive cells in each region of interest was counted automatically by calculating the mean image pixel intensity and applying a threshold, with validation by microscopic counting.

#### Quantification

To quantify activated excitatory or inhibitory neurons in rat cortex, 40 μm-thick transverse sections of were immunolabeled with c-Fos, PV, CamKII antibodies and counted the number of double positive cells (activated excitatory neurons: c-Fos and CamKII, activated inhibitory neurons: c-Fos and PV). At least 4 brains were analyzed for each group. All quantifications in images were analyzed in ImageJ software.

### Statistical analysis

The study data were analyzed using OriginPro version 9.1 software (OriginLab, Northampton, MA, USA). The data were assessed for normality using the Kolmogorov-Smirnov test. The numbers of cells expressing c-Fos were then compared between the study groups using one-way analyses of variance. The Bonferroni post-hoc test was used to detect significant differences between groups for each region of interest and specific cell type. If no significant differences were detected, Kruskal-Wallis one-way analysis of variance by ranks with Dunn’s method was used to compare the specific numbers of cells co-labeled with c-Fos between the study groups and for post hoc comparison. A *p*-value <0.05 was considered statistically significant.

### Computational simulation

#### Stimulus-induced potential field

We constructed a three-dimensional finite element model of a rat head using computed tomography images of a rat. The rat brain imaging was performed using a volumetric micro-CT scanner (NFR Polaris G90C; NanoFocusRay, Ikson, Korea). The image size was 1024×1024 pixels with 434 slices and the voxel size was 0.0698×0.0698×0.1396 mm^3^. Manual segmentation was performed using Seg3d to guarantee continuity and to improve the accuracy of segmentation. The model consisted of the scalp, skull, and cerebrospinal fluid (CSF); we included the brain after shrinking the CSF layer by 5 mm. The electrodes were modeled in accordance with the surgical procedures. The rat head model was generated by an optimized tetrahedral mesh using Iso2Mesh toolbox [30], TetGen [31], and MATLAB.

The electrical properties of each tissue, taken from averaged human conductivity values, were assigned as follows (S/m): skin, 0.45; skull, 0.01; CSF, 1.65; brain, 0.2; and electrode, 5.5e07. The potential field were calculated by solving the quasi-static Laplace equation via COMSOL Multiphysics (v5.3, COMSOL. Inc., Burlington, MA, USA) using the finite element method. We applied the conjugate gradient method with preconditioning of an algebraic multigrid (relative tolerance, 1×10^-6^; Fig. 2)

**Figure 2.**
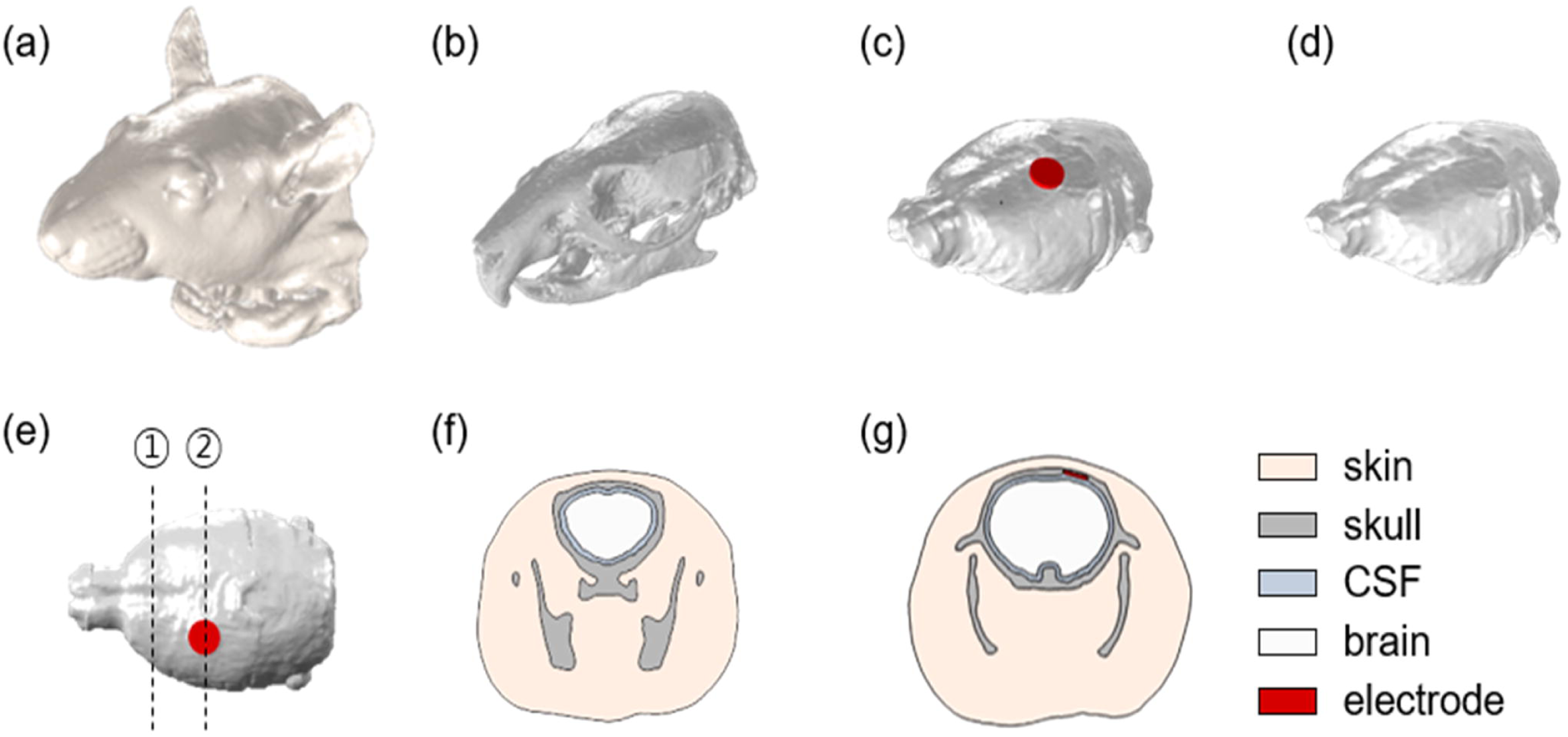
High-resolution computed tomography scan based on an anatomically realistic rat head model. The anatomically realistic rat head model consisted of four layers of skin (A), skull (B), cerebrospinal fluid (C), and brain (D). The electrodes are placed in accordance with the coordinates used in the experiment (E), and a cross-section, following the black dotted line shown in (E), passing the reference (F) and the active electrode (G) are shown.

#### Neuronal responses to the electrical field

Single-compartment models with Hodgkin-Huxley properties were modified to represent regular spiking excitatory neurons and fast spiking interneurons [22]. The constants and parameters were unchanged from the original models, which resulted in different firing patterns with respect to various frequencies and the strength of sinusoidal stimulation. The neuronal models were implemented in NEURON [32].

To simulate neuronal responses according to the predicted potential distributions calculated in the rat head model, we constructed a multi-scale model that virtually combined the single-compartment neuronal models with the rat head model [33, 34]. The multi-scale model consisted of the following two-step process. First, the excitatory and inhibitory neurons were distributed on the cortical surface. Second, the external stimulus calculated using the rat head model was applied to neuronal models. Therefore, the external input I(t) = I_0_*f*(*t*) was added to the cable model where I = ∂^2^V/∂*s*^2^ approximates the amplitude of the stimulus-induced transmembrane current and *f*(*t*) represents the pulse waveform. The amplitude of the extra current I_0_ is determined by an “activating function” that evokes activation of neuronal models [35]. S is the direction that is locally parallel with the fiber, and we assumed the fiber direction to be the normal direction of the close element comprising the cortical surface. The I_0_ was calculated at each neuron’s position in the rat head model using COMSOL with MATLAB. For the waveform, we simulated continuous sinusoidal stimulation at 20, 40, 60, 80, and 100 Hz in accordance with the animal experiment.

## RESULTS

### Quantification of neuronal activity from expression of c-Fos after SEBS

c-Fos is an immediate-early gene that responds transiently and rapidly to various stimuli and is a good marker of neuronal activation in the brain [36]. To identify neuronal activation by SEBS in the rat brain, we performed SEBS at various frequencies (20, 40, 60, and 100 Hz) 24 h/day for 1 week (Fig. 1A). The immunoreactivity of c-Fos shows the neuronal activation density map at two different bregma levels after SEBS at 20, 40, 60, and 100 Hz in the experimental groups and in the sham group (Fig. 3A). Activation of c-Fos by SEBS was propagated to various brain regions including not only the motor and sensory cortices but also the deeper brain, including the striatum and thalamus. To elucidate the neuronal activation in the various regions, we performed automated c-Fos positive cell counts in four regions of interest (motor cortex, sensory cortex, striatum, thalamus) among the groups (Table S1). SEBS at 40 Hz resulted in the highest significant increment of c-Fos-positive cell count in the contralateral sensory cortex (*p*<0.05). In the 20 Hz SEBS group, activation was highest in the thalamus (*p*<0.001). Regardless of frequency, the increments in neuronal activity were significantly greater in the EBS groups than in the sham group (Fig. 3B, Table S1).

**Fig. 3.**
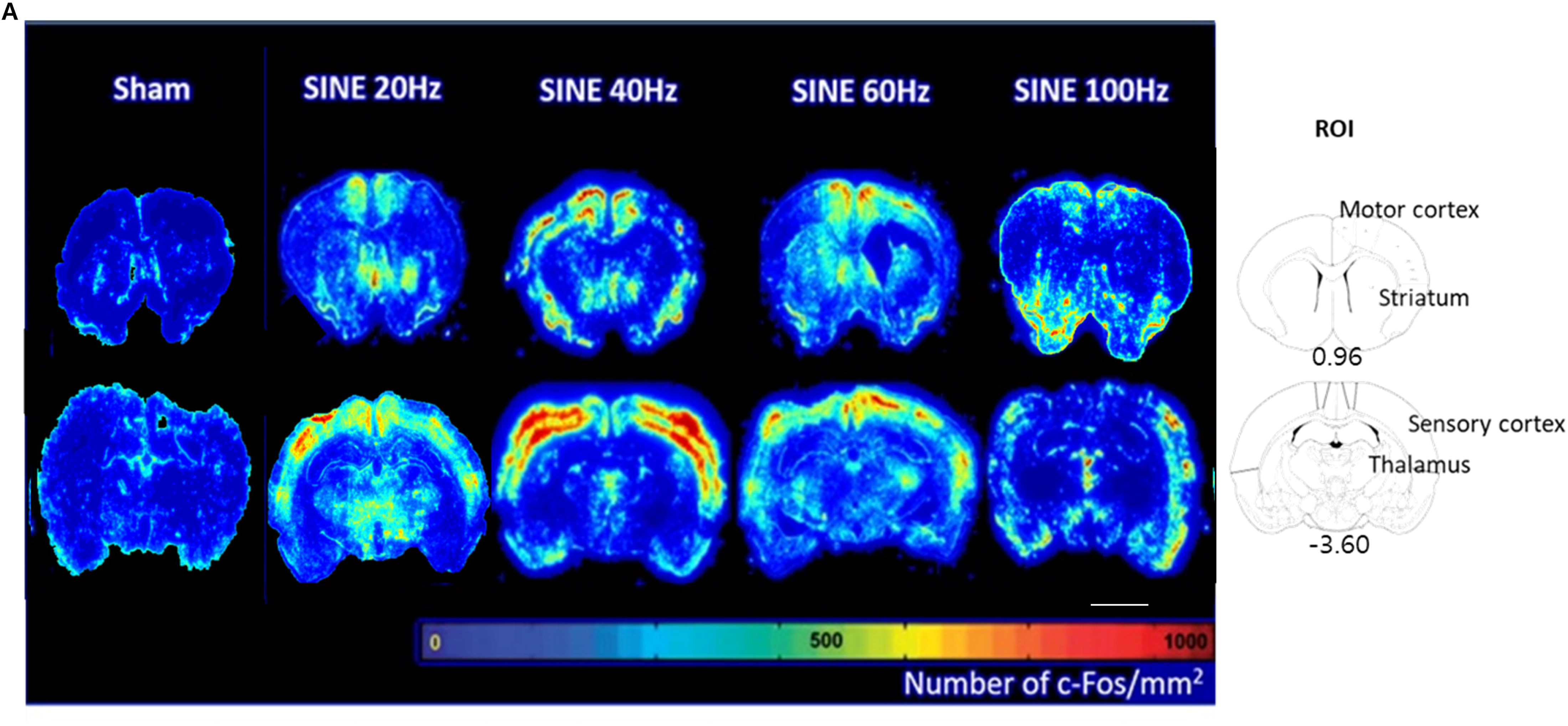
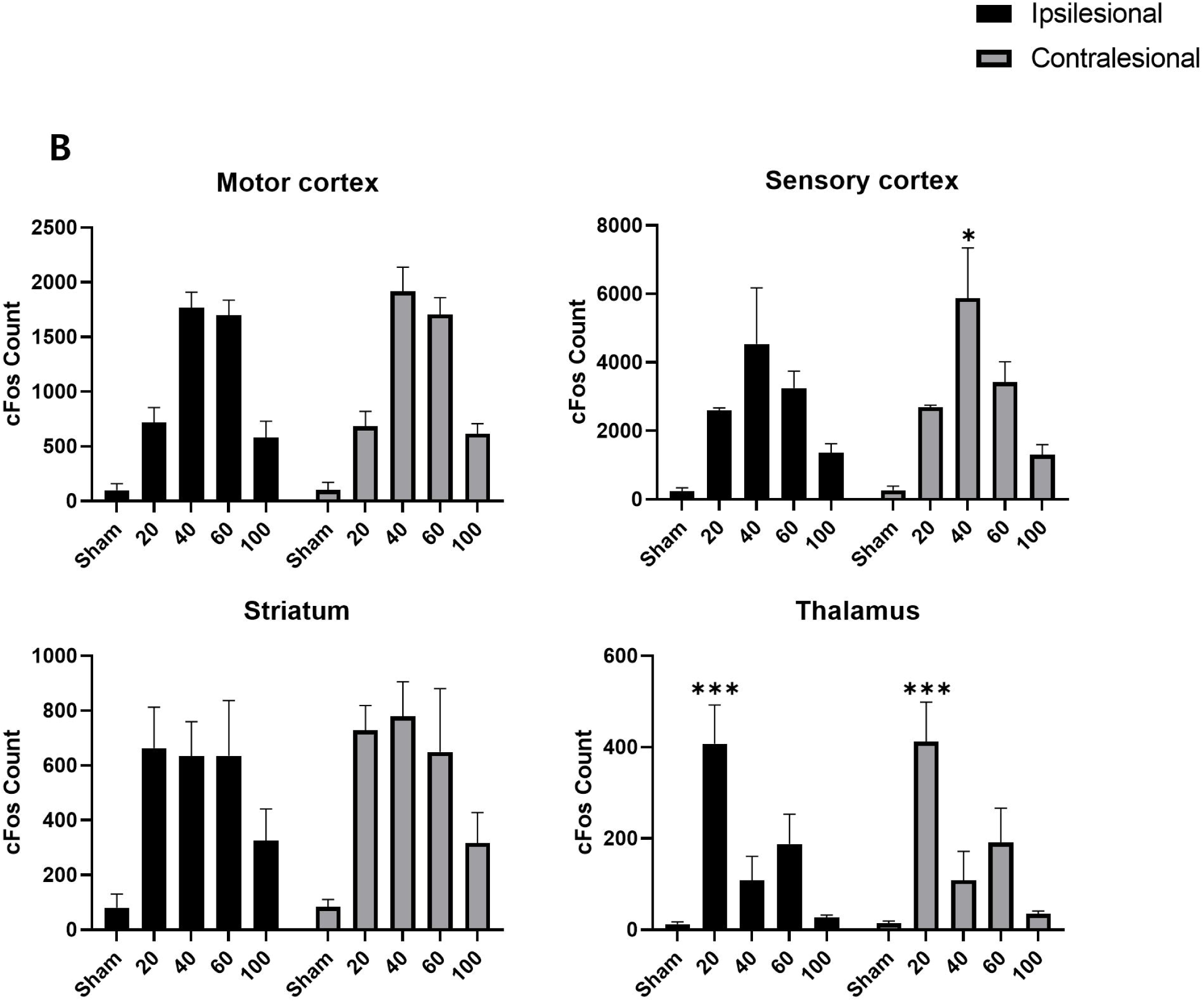
Expression of c-Fos after sensory-parietal cortical stimulation. (A) Cell-density maps for c-Fos-positive cells at three bregma levels (+0.96, −3.60) in the sham operation group and the experimental groups that received SEBS at 20, 40, 60, or 100 Hz (left), together with the atlas reference section (right). (B) Automated cell counts in four regions of interest, i.e., the motor cortex, sensory cortex, striatum, and thalamus. Comparing stimulated groups with the sham group showed that SEBS increased cFos activity. SEBS at 40 Hz achieved the highest increment in c-Fos in the motor and sensory cortex and SEBS at 20 Hz showed the highest increment of c-Fos in the thalamus. The error bars represent the standard error of the mean. \**p*<0.05 and \*\*\**p*<0.001 compared with the other study groups, one-way analysis of variance with Bonferroni post-hoc test. Scale bar: C, 3 mm. EBS, electrical brain stimulation; n.s., not statistically significant

### Quantification of sensory cortical cell type-specific activity using co-labeling with c-Fos expression after SEBS

Excitatory/inhibitory balance is required for correct functioning of the brain. Because SEBS modulates the excitatory/inhibitory neuronal balance, it has been applied as effective treatment for various neurological disorders [37–39]. To elucidate the functional mechanism of SEBS, we investigated the types of neuronal cells that are activated after SEBS. Immunohistochemistry was performed with calmodulin-dependent protein kinase II (CaMKII) antibody for excitatory neurons and PV antibody for inhibitory neurons. Activated inhibitory neurons (PV^+^, c-Fos^+^) and excitatory neurons (CaMKII^+^, c-Fos^+^) were analyzed (Fig. 4).

**Fig. 4.**
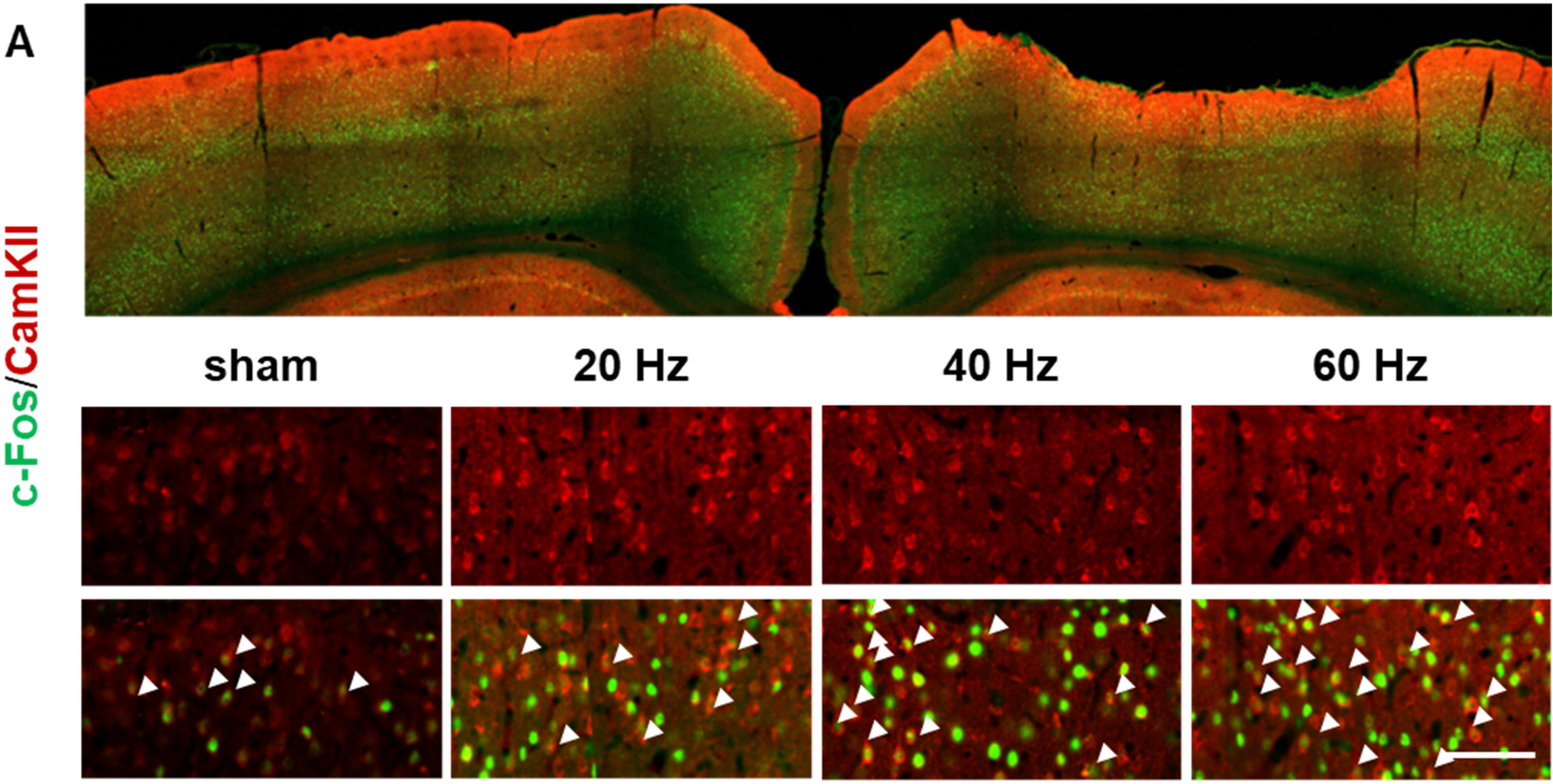
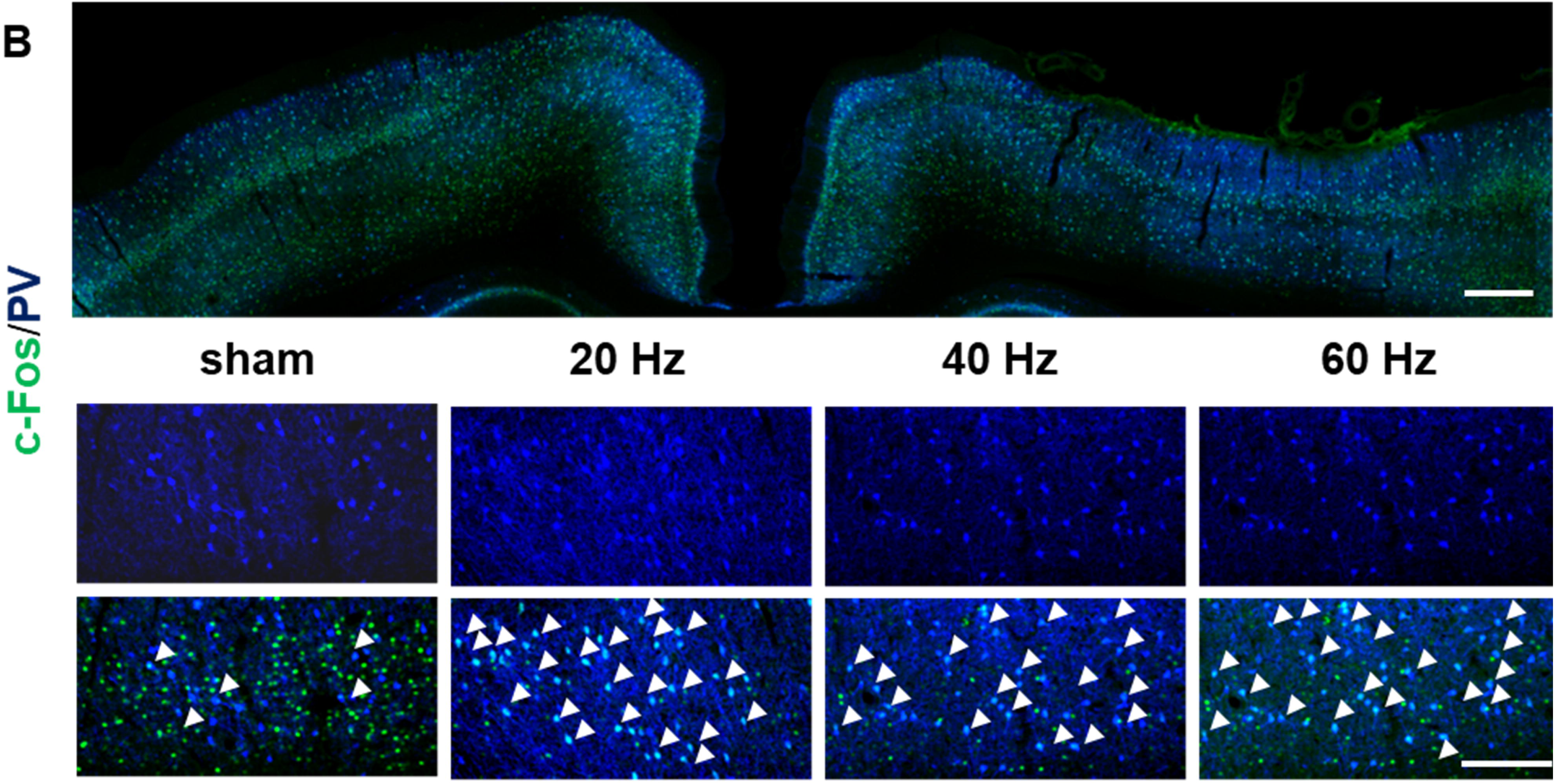
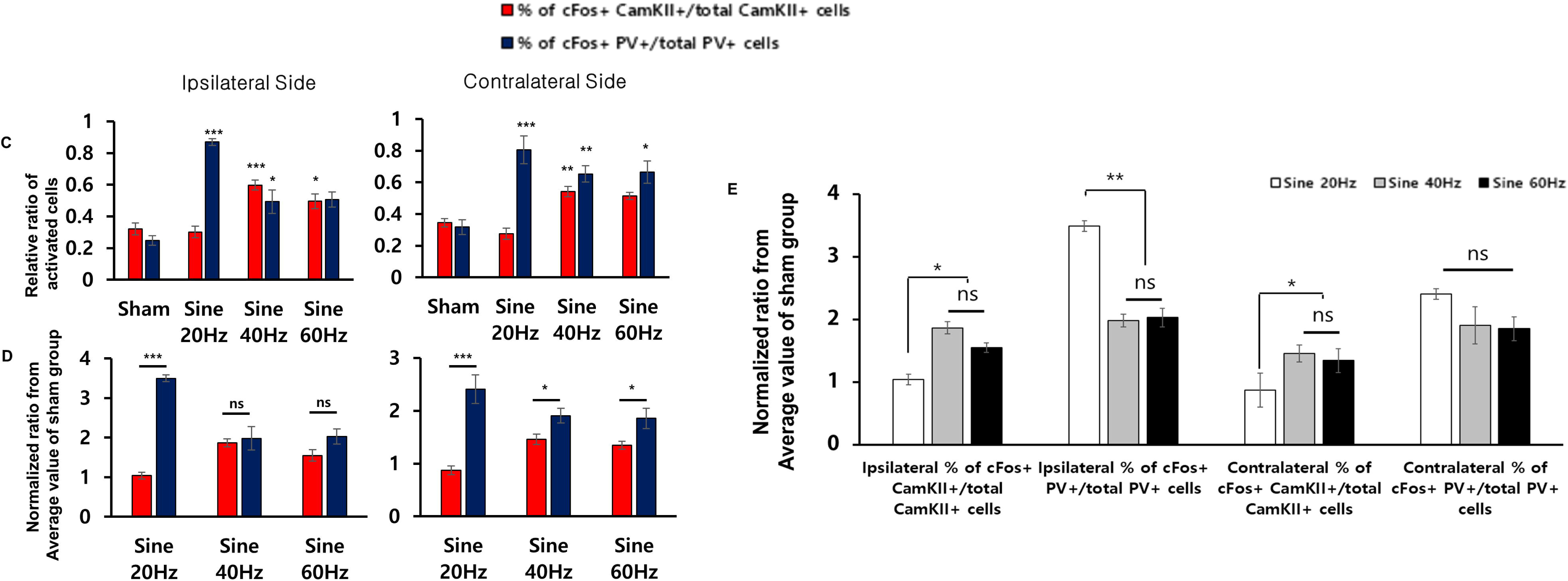
Histological confirmation of differential electrical stimulation and pattern of effect during cortical stimulation of the sensory parietal cortex for 1 week. (A) Immunostaining of a rat cortex with CaMKII and c-Fos antibodies. CaMKII and c-Fos double-positive cells indicate activated excitatory neurons. (B) Immunostaining of a rat cortex with PV and c-Fos antibodies. PV and c-Fos double-positive cells indicate activated inhibitory neurons. (C) Quantification of the activated neuron ratio in the rat cortex in response to sinusoidal electrical stimulation at different frequencies, i.e., sham, 20 Hz, 40 Hz, and 60 Hz. (D, E) Quantification of the ratio of activated neurons from the mean value for the rat cortex in the sham group in response to sinusoidal electrical stimulation at frequencies of 20, 40, and 60 Hz. (D) Comparison between CaMKII and PV at specific frequencies. (E) Comparison between different frequencies according to specific regional cell type. The white arrow represents c-Fos and CaMKII or c-Fos and PV double-positive cells. The error bars represent the standard error of the mean. \**p*<0.05 and \*\*\**p*<0.001 compared with the control, one-way analysis of variance with Bonferroni post-hoc test. Scale bars: B, 500 μm and 50 μm. CaMKII, calmodulin-dependent protein kinase II; PV, parvalbumin; n.s., not statistically significant

20Hz SEBS markedly improved the activity of inhibitory neurons in both cortices but not that of excitatory neurons when compared with the sham group (ipsilateral side; 86.98±2.12%, *p*<0.001; contralateral side; 80.54±8.88%, *p*<0.001). Even though 40Hz and 60Hz SEBS increased the activity of inhibitory neurons, excitatory neurons were more activated than those in the sham group (40 Hz on ipsilateral side, 59.82±3.25%, *p*<0.001; 40 Hz on contralateral side, 54.22±3.29%, *p*<0.01; 60 Hz on ipsilateral side, 49.54±4.70%, *p*<0.05; 60 Hz on contralateral side, 51.30±2.26%, *p=*0.09; Fig. 4C).

We also calculated the normalized ratio from the average value in the sham group to compare the excitatory and inhibitory neurons at each frequency (Fig. 4D) and specific regional neurons between three different frequencies (Fig. 4E). The activity of inhibitory neurons was markedly increased by 20Hz SEBS when compared with the activity of excitatory neurons (ipsilateral side, 3.49±0.09 and 1.04±0.08, respectively, *p*<0.001; contralateral side, 2.41±0.27 and 0.87±0.08, *p*<0.001). The activity of inhibitory neurons was increased by 40Hz and 60Hz SEBS; however, the increase was only statistically significant on the contralateral side (40 Hz on ipsilateral side, 1.98±0.29 and 1.87±0.10; 40Hz on contralateral side, 1.91±0.14 and 1.46±0.09, *p*<0.05; 60 Hz on ipsilateral side, 2.03±0.19 and 1.55±0.15; 60Hz on contralateral side, 1.86±0.19 and 1.35±0.08, *p*<0.05; Fig. 4D).

The activity of excitatory neurons that received 40Hz and 60Hz SEBS was significantly greater than that of excitatory neurons that received 20Hz SEBS on both the ipsilateral and contralateral sides (*p*<0.05 and *p*<0.05, respectively); however, the activity of inhibitory neurons was significantly greater in response to 20Hz SEBS than in response to 40Hz and 60Hz SEBS only on the ipsilateral side (*p*<0.01; Fig. 4E)

CaMKII-positive excitatory neurons were predominantly located in layers 2/3 and 6 but PV-positive inhibitory neurons were mainly found in layers 4 and 5 [40, 41]. We quantified the number of neurons activated according to the cortical layer in which they were located by dividing the cortex into 10 bins (Supplementary Fig. 2A, 2B). We found that 40Hz SEBS activated excitatory neurons in the whole cortical layer but that 60Hz SEBS only activated neurons in layers 2/3 and 6. Although 20Hz SEBS did not activate excitatory neurons, it activated inhibitory neurons in the entire cortical layer, and 40-Hz and 60-Hz SEBS activated inhibitory neurons in the deeper layer (Supplementary Fig. 2C, 2D).

### Computational simulation for observing the effect of SEBS on geometrical and neuronal responses at different frequencies

We performed computational simulation to identify the geometrical impact of SEBS on activation of neurons and their firing rate. First, the current density induced by a 1-V stimulus amplitude was computed (Fig. 5). As expected, there was a higher current density in the CSF because of higher conductivity, and the current density was strongest in the brain area directly beneath the electrode. A high current density is observed at the edge of the active electrode because of the edge effect whereas the reference electrode has less of an edge effect because it is smaller. Second, in order to elucidate the functional mechanism of different excitatory/inhibitory neuronal activation by SEBS, we simulated single neuron responses by increasing the external stimulus. The action potentials (APs) efficiency, which is the percentage of action potentials per stimulation pulse, is shown in Fig. 6, and clearly shows the firing patterns. Generally, there was no firing for a lower stimulus amplitude with a higher stimulus frequency and there was burst firing for a stronger stimulus amplitude with a lower stimulus frequency. Following current-controlled stimulation, phase-locked firing patterns were frequently observed in both inhibitory and excitatory neuron models. The stimulus amplitude needed to evoke APs was lower for inhibitory neurons than for excitatory neurons. Third, we coupled individual neurons to spatial patterns of a stimulus-induced current field calculated using a rat head model (Fig. 7). Consistent with the spatial distributions of current density shown in Fig. 5, the neurons were activated directly beneath the electrodes regardless of stimulus frequency and type of neuronal model. Inhibitory neurons were more strongly activated than excitatory neurons because of intrinsic differences between these two types of neurons. The stimulus amplitude needed to evoke neuronal activation increased monotonically with stimulus frequency; therefore, 20-Hz SEBS induced the strongest activation in both inhibitory and excitatory neurons, with shrinking of the activated area as the frequency increased (Fig. 7).

**Fig. 5.**
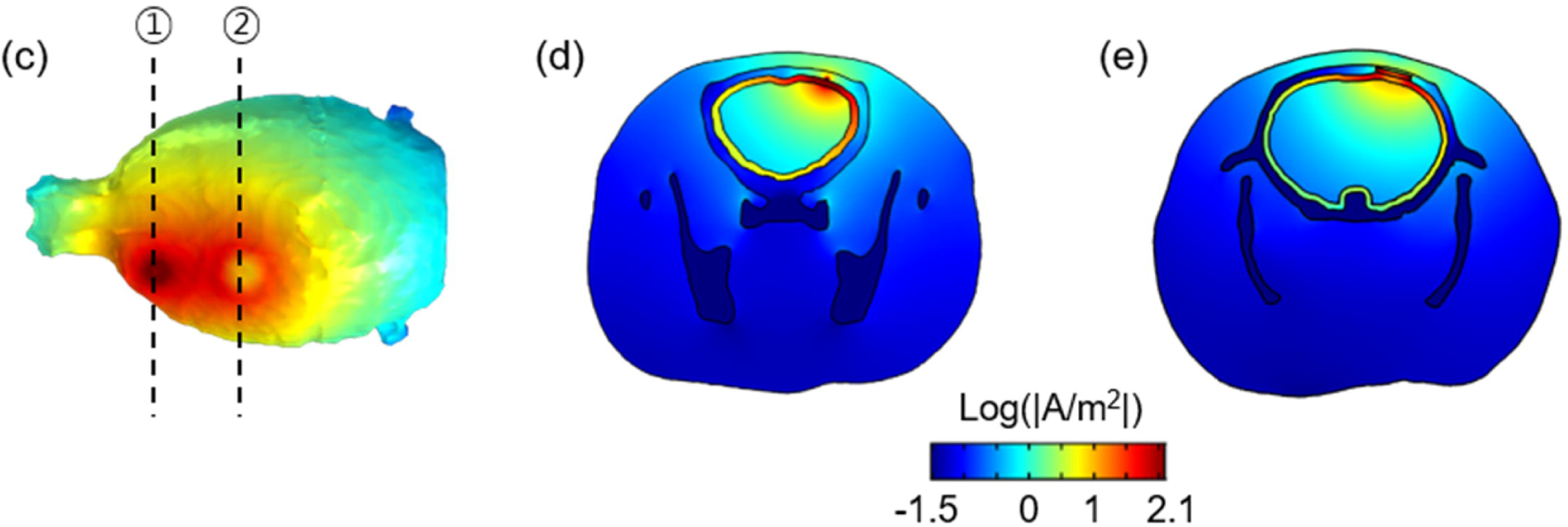
Simulated current density distribution. The spatial distributions of current density induced by 1V stimulus amplitude are visualized at the surface of the brain (A) and the cross-section passing the reference (B) and active (C) electrodes (following black dotted line shown in (A)) are shown.

**Fig. 6.**
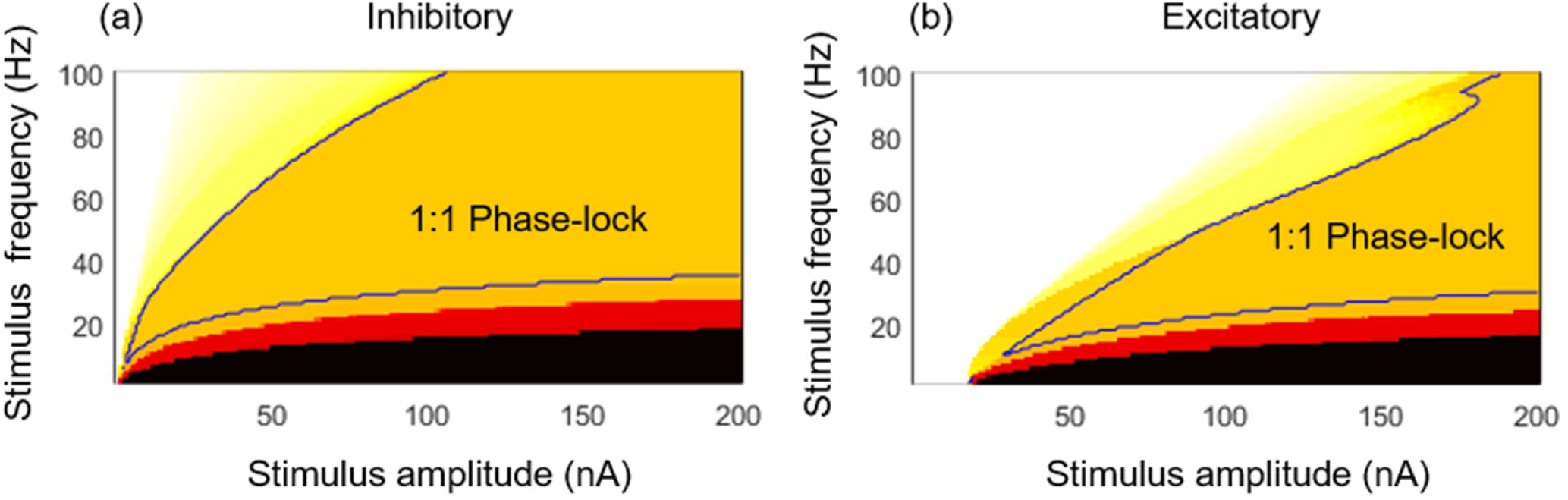
Map showing the relationship between firing frequency and sinusoidal stimulation. The spatial distributions of the firing rate for the inhibitory (A) and excitatory (B) neuron model induced by different stimulus amplitudes and stimulus frequencies are depicted. The firing rate is analyzed by action potentials (APs) efficiency, which define percentage action potentials per stimulation pulse. An APs efficiency value >1 indicates burst, a value of 1 indicate phase lock, and a value of <1 indicates intermittent firing behavior. The blue contour lines represent the 1:1 phase-locked firing region.

**Fig. 7.**
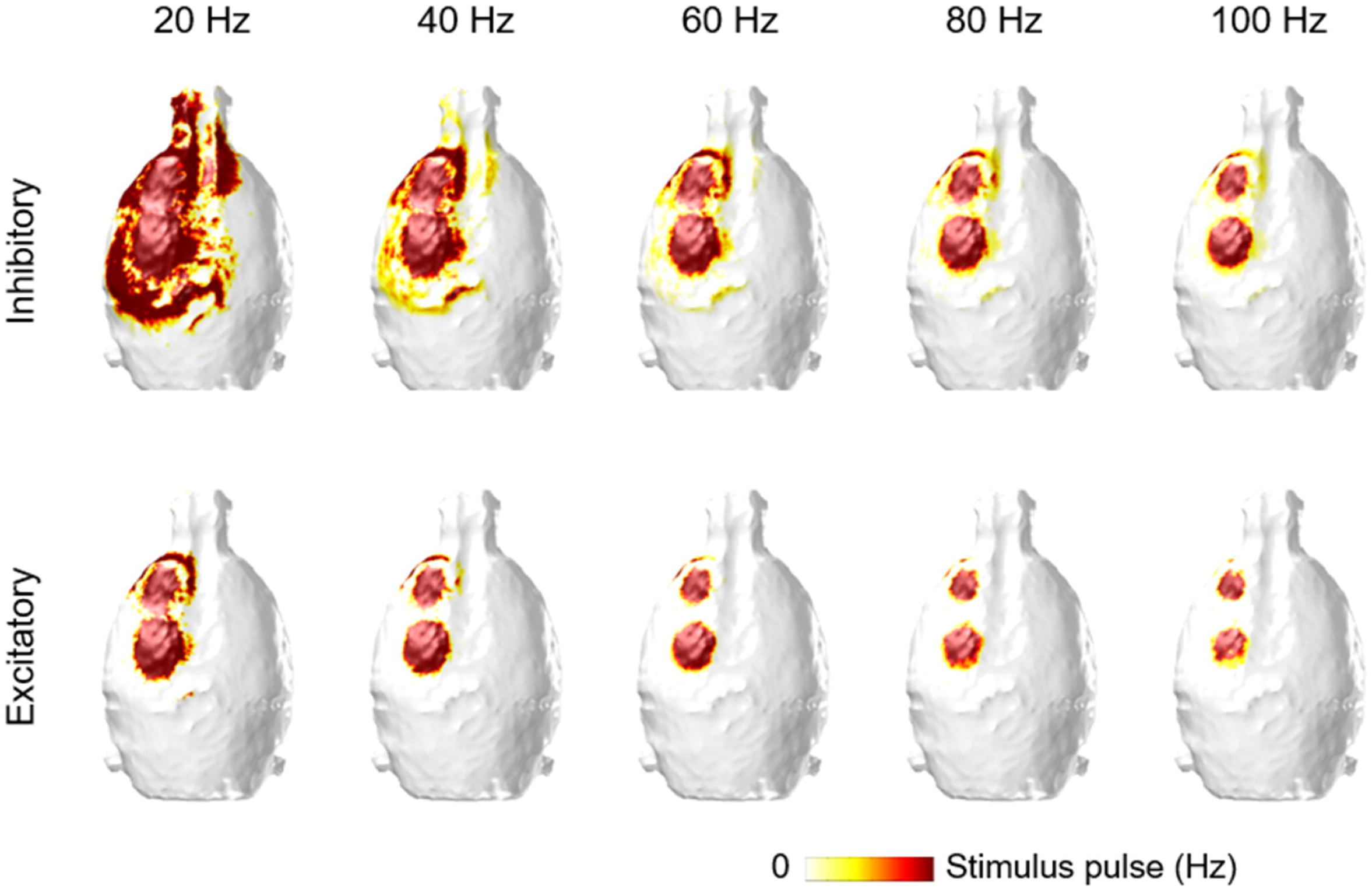
Simulated spatial distribution of firing rate. The spatial distributions of the firing rate induced by a 1-V stimulus amplitude are visualized on the surface of the brain for excitatory and inhibitory neurons by increasing the stimulus frequency in steps of 20 Hz.

## DISCUSSION

The precise mechanism underlying neuronal activation by SEBS is not well understood. Electrical stimulation of the sensory and motor cortex usually focus on selective stimulation of cortical pyramidal cells because pyramidal neurons are known to be the primary activators of the corticospinal tract and may provide the main input to the direct pathway [42, 43]. Based on experimental results showing higher activation in inhibitory neurons [15, 44], the possibility of cell type-specific individual neuron responses being a bridge to interpolation of neural networks has been raised.

To improve our understanding of this mechanism, we performed an animal experiment and biocomputation. Synaptic connectivity and the strength of individual neurons are usually assessed by intracellular recordings; however, it is impractical to record data for all neurons within a neural circuit. As an alternative, we observed the firing properties of the neural circuit by eliciting a response in individual pyramidal cells and interneurons displaying a diverse activation pattern, reflecting their anatomical structure using a computational study.

### Neuronal responses in vivo

Our experimental study revealed expression of c-Fos, which represents neuronal activity [36], to be higher in all of the SEBS groups than in the sham group. Cell type-specific analysis showed that neuronal activity was stronger in PV+ interneurons, regardless of type of SEBS, than in the sham group. This is consistent with a previous result for transcranial direct current stimulation (tDCS), which predominantly modulates interneurons [45]. Interneurons are primarily inhibitory in the central nervous system and their main role is to conduct flow of neuronal signals between a motor neuron or sensory neuron in a neural circuit. PV interneurons are crucial when performing high-order functions such as learning and decision-making and also regulate the activity of pyramidal neurons [46]. Therefore, our findings afford a clue for understanding the effect of current SEBS, such as tACS, in specific neuronal diseases.

SEBS of the sensory parietal cortex and motor cortex enhanced neural activity locally beneath the area of the electrical field, unlike at the striatum and thalamus, which were distant from the electromagnetic field used in our experiments. SEBS differentially affects the local electromagnetic strength at the circuit level. Thalamic activation was greater at 20 Hz than at other frequencies. These findings suggest that a neural circuit, e.g., the cortico-striatal-thalamic circuit of the salience network or the corticothalamic circuit, will also be influenced by SEBS. This modulation of circuitry could represent additional clue for therapeutic intervention.

The effects of EBS were divided into those that occurred during stimulation and those that occurred after stimulation. Those that occur during stimulation are solely dependent on changes in the membrane potential while those that occur after stimulation depend on membrane depolarization [15] and synaptic modulation [47]. The aftereffects of cathodal tDCS depend on modulation of glutamatergic synapses [15]. The mechanism of action of SEBS in the context of specific neurons could be different from the simple summation of anodal and cathodal tDCS effects The phenotypic effect is represented as the balance of excitatory and inhibitory neuronal responses. Our simulation results showed that 20Hz SEBS evokes the strongest inhibitory and excitatory neuronal responses in both model of rat and human brain (Fig. 7, Supplementary Fig. 1). Since the inhibitory neuronal simulation responses are much larger than excitatory simulation neuronal responses in the group that received 20Hz SEBS, our clinical findings can be interpreted as meaning that stronger inhibitory neuronal responses occurred in the group that received 20Hz SEBS (Fig. 4D) and that excitatory neuronal activity occurred in the groups that received 40Hz or 60Hz SEBS (Fig. 4E).

### Individual neuronal responses in silico

We simulated the activation of an inhibitory and excitatory neuron model by extracellular stimulation and examined the relationship between the neuronal firing profile and stimulus frequency in respect to realistic stimulus-induced field distributions. We found neuronal excitability to be reduced in response to a strong stimulus frequency and inhibitory neurons were more sensitive than excitatory neurons to sinusoidal stimulation.

We adapted an established model of excitatory and inhibitory neurons from Mahmud et al [22]. In their model, an applied extracellular stimulation current was calculated by a derivative of potential field, which can be interpreted by intracellular current stimulation. In our study, as an alternative, we adjusted the stimulus amplitude via the activating function calculated using the simulated current field in the rat head model. Therefore, we were able to take into account the neuron’s location relative to the electrodes, which may be an important determinant of neuronal polarization [48].

Additionaly we simulated the spatial distribution of the firing rate using a human head model to investigate the impact of the complex geometry of the human brain on neuronal activation, as depicted in Supplementary Fig. S1. We used a previously developed human head model [48, 49] and coupled it to the same type of neuronal model in the rat head. As we placed the reference electrode on the chest far from the active electrode for the human model, the activation of neurons was restricted to the area of the sensory cortex directly beneath the electrode. The complex patterns reflecting the complex anatomy of the brain were observed to result in activations in the sulcal wall. As expected, a 20-Hz sinusoidal stimulation produced the largest areas of activation and the stimulus threshold was lower for inhibitory neurons than for excitatory neurons (Fig. 7 and Supplementary Fig. S1).

Most computational approaches have presented neuronal excitability with sinusoidal stimulation subject to a uniform electric field. Aspart et al reported a frequency-dependent polarization profile using a biophysically detailed model of pyramidal neurons in response to a weak uniform electric field [50]. Yi et al suggested that the geometric features of a neuron model play a crucial role in determining the polarization when using a two-compartment neuron model with a uniform electric field [51]. Thus, in the future, we need to achieve morphological features of neuronal models for precise simulation; however, while most simulation studies investigating neuronal activation involve intracellular or extracellular stimulation with a uniform electric field, we propose a multi-scale model and thus we could consider extracellular field induced in the brain by the stimulating electrodes. This study has some limitations. First, c-Fos activation reflects rapid responses after SEBS. In this study, SEBS was performed for 1 week, so measurement of c-Fos cannot fully represent the neuronal responses that occur with long-term SEBS. However, 1 week of SEBS can modulate synapses at the neuronal circuit level and we assume that our c-Fos density mapping shows the chronic effects of SEBS. Second, the translation of individual neuronal responses into oscillations at the network level is not trivial. However, modulation of individual neurons may provide evidence of entrainment of neural oscillations at the network level through which neurons are recruited by sinusoidal stimulation. Future studies should incorporate synaptic connections, which may allow more precise and effective application of epidural EBS in both the clinical and basic research settings.

## CONCLUSION

We examined the change in stimulation parameters when stimulating brain cells and the quantity of each component of brain cells to determine which cells are differentially influenced after delivering cortical stimulation. Our findings were derived from in vivo and in silico experiments and analyzed in an integrated manner. We found that 20-Hz SEBS is the most effective frequency for selective inhibitory cortical stimulation and that 40-Hz SEBS is the most effective frequency for selective excitatory cortical stimulation. We assumed the mechanism involves computational simulation whereby 20-Hz SEBS differentially stimulates inhibitory neurons and inhibits excitatory neurons sequentially. In order to obtain a functional effect by applying SEBS clinically, it is necessary to consider the effect of SEBS on specific neurons and neuronal circuits in specific neurological disorders.

## Supporting information

supplementary_result

highlights

## Acknowledgments

The authors are grateful to Professor MC Lee for his enthusiastic discussion about differential cell counting based on his neuropathologic knowledge.

## Data statement

The datasets generated and analyzed during the current study are available from the corresponding author on reasonable request.

## FIGURE LEGENDS

**Figure S1.**
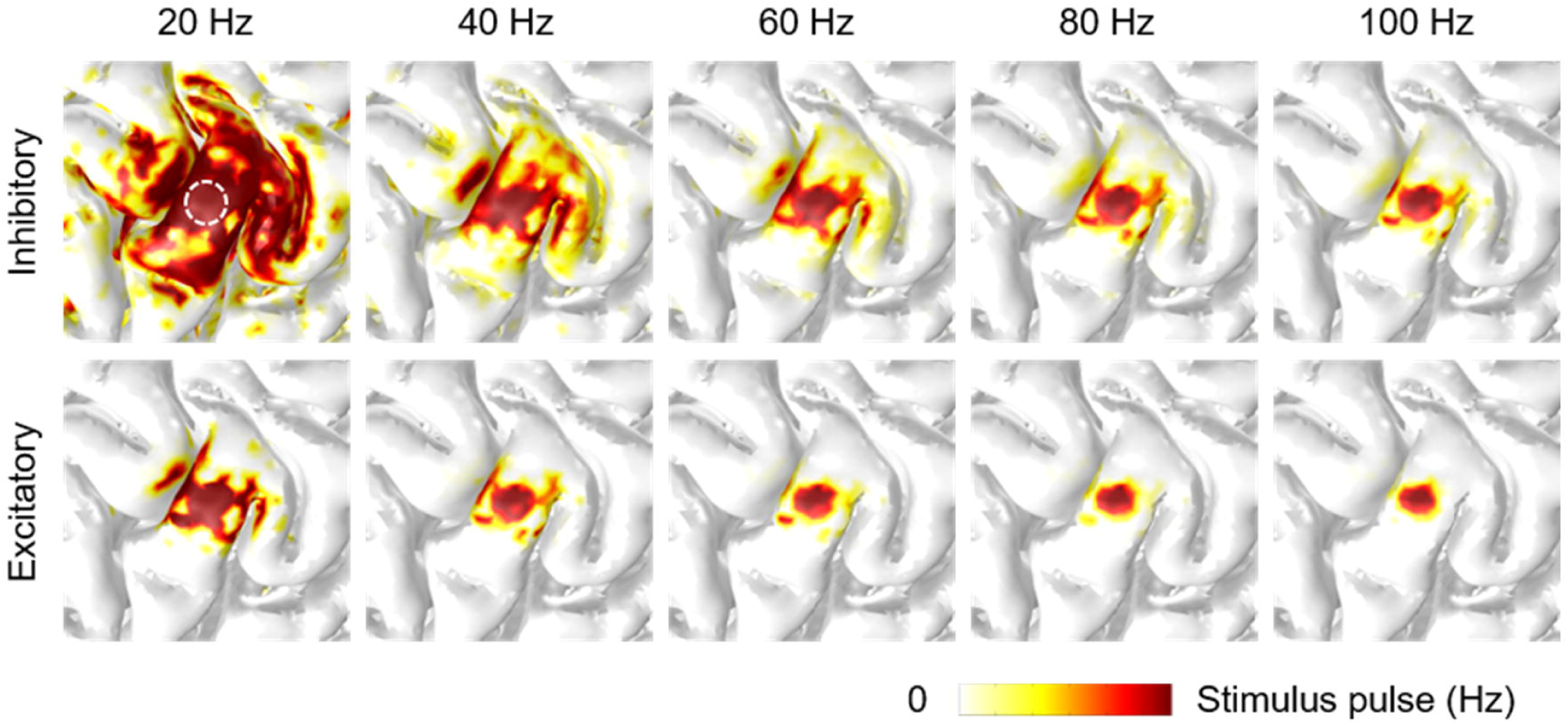
Simulated spatial distribution of firing rate subject to the human model. Simulated spatial distribution of firing rate subject to the human model. The spatial distribution of firing rate induced by 1V stimulus amplitude are depicted at the cortical surface for excitatory and inhibitory neurons, by increasing stimulus frequency in steps of 20 Hz. The active electrode targeting sensory cortex is marked by while-colored and dotted line in 20 Hz stimulus-induced inhibitory neuronal responses.

**Figure S2.**
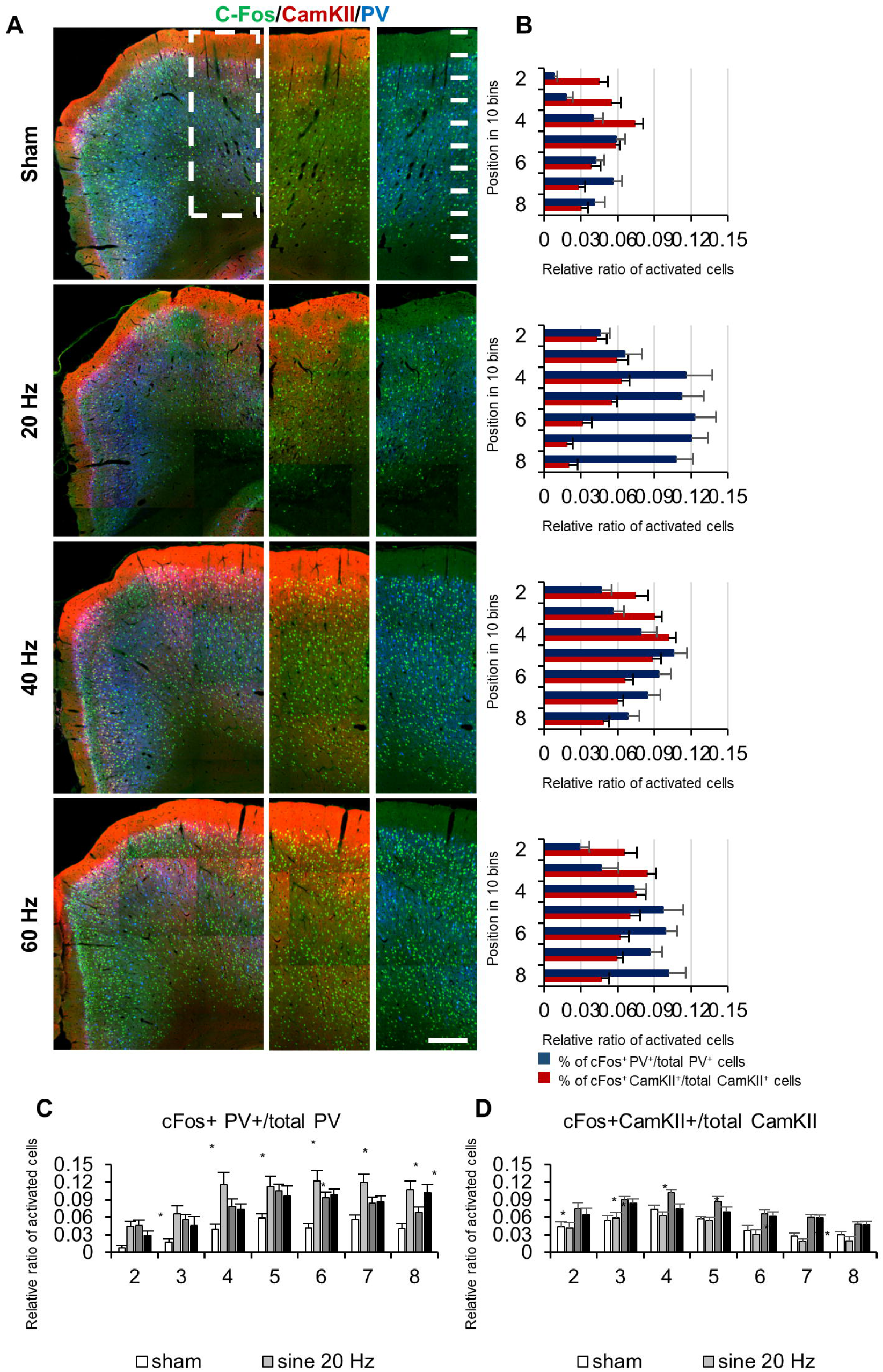
Investigation of activated neuronal cell types at the cortical layer localization. **(A)** Representative image of CamKII, PV and c-Fos immunostaining at rat cortex. **(B-D)** Distribution analysis of activated neurons in the rat cortex. Error bars represent S.E.M. *p < 0.05 compared with sham control, two way analysis of variance on ranks with Dunn’s method. Scale bars: B, 200 µm

**Table S1.** The number of c-Fos expression after sensory-parietal cortical stimulation. Automated cell counts of sham, 20, 40, 60, 100Hz group in four different regions of interest: motor cortex, sensory cortex, striatum, thalamus. Comparing stimulated groups with sham operation group showed that sinusoidal EBS increased c-Fos activity.

|  | Ipsilateral |  |  |  |  | Contralateral |  |  |  |  |
| --- | --- | --- | --- | --- | --- | --- | --- | --- | --- | --- |
| unit: cells | Sham (n=5) | 20 Hz (n=4) | 40 Hz (n=7) | 60 Hz (n=4) | 100 Hz (n=7) | Sham (n=5) | 20 Hz (n=4) | 40 Hz (n=7) | 60 Hz (n=4) | 100 Hz (n=7) |
| Motor cortex | 98.4 ± 24.14 | 717 ± 59.85 | 1768.29 ± 49.61 | 1697 ± 60.72 | 577.86 ± 53.61 | 101.8 ± 28.06 | 683.75 ± 59.20 | 1916.57 ± 78.04 | 1706.25 ± 66.38 | 616 ± 32.10 |
| Sensory cortex | 245 ± 82.61 | 2595.25 ± 33.78 | 4539 ± 573.03 | 3248.5 ± 214.85 | 1554.14 ± 129.79 | 264.4 ± 47.28 | 2687 ± 27.17 | 5877.14 ± 513.63 | 3423.75 ± 256.16 | 1530.29 ± 124.85 |
| Striatum | 80.2 ± 44.86 | 662.5 ± 65.23 | 634.43 ± 43.77 | 635 ± 87.39 | 326.29 ± 40.27 | 83.6 ± 10.59 | 727.5 ± 39.58 | 779 ± 44.31 | 648.75 ± 100.24 | 316.71 ± 39.04 |
| Thalamus | 12.4 ± 4.45 | 407.25 ± 36.79 | 108.29 ± 18.31 | 187.75 ± 28.31 | 26.86 ± 1.94 | 14 ± 2.12 | 412 ± 37.60 | 108.57 ± 22.05 | 191.25 ± 32.55 | 35 ± 2.26 |

