## supplementary_result for "Effect of sinusoidal electrical cortical stimulation on brain cells"

Seungjun Ryu^1†^, Kyung-Tai Kim^2†^, Hyun Seo^3†^, RaGyung Kim^1^, Jongwook Cho^1^, Jiyoung Park^1^, Sunwoo Lee^1^, Hanlim Song^1^, Hyoung-Ihl Kim^1*^

^1^Department of Biomedical and Science and Engineering (BMSE), Institute of Integrated Technology (IIT), Gwangju Institute of Science and Technology (GIST), Gwangju 61005, Korea

^2^ Jeonbuk Department of Inhalation Research, Korea Institute of Toxicology, 30 Baekhak1-gil, Jeongeup, Jeollabuk-do 56212, Republic of Korea

^3^School of Electrical Engineering and Computer Science, Gwangju Institute of Science and Technology (GIST), Gwangju 61005, Korea

*Corresponding author.

**Supplementary Data**

**1. Supplementary Methods**

### Statistical analysis

To quantify activated excitatory or inhibitory neurons in the rat cortex, cortex was divided into 10 bins. Of the 10 bins, cerebral gray matter were located 2 to 8 bins. Activated excitatory neurons (c-Fos^+^, CamKII^+^ neurons) and activated inhibitory neurons (c-Fos^+^, PV^+^ neruons) were quantified.and two-way analyses of variance were used to compare the number of activated neurons using statistical analysis software (SigmaStat,3.5, Systat Software Inc.). The statistical significance was set at *p* < 0.05.

**2. Supplementary Results**

**Table S1. The number of c-Fos expression after sensory-parietal cortical stimulation.** Automated cell counts of sham, 20, 40, 60, 100Hz group in four different regions of interest: motor cortex, sensory cortex, striatum, thalamus. Comparing stimulated groups with sham operation group showed that sinusoidal EBS increased c-Fos activity.

**Figure S1. Simulated spatial distribution of firing rate subject to the human model.** Simulated spatial distribution of firing rate subject to the human model. The spatial distribution of firing rate induced by 1V stimulus amplitude are depicted at the cortical surface for excitatory and inhibitory neurons, by increasing stimulus frequency in steps of 20 Hz. The active electrode targeting sensory cortex is marked by while-colored and dotted line in 20 Hz stimulus-induced inhibitory neuronal responses.

**Figure S2. Investigation of activated neuronal cell types at the cortical layer localization. (A)** Representative image of CamKII, PV and c-Fos immunostaining at rat cortex. **(B-D)** Distribution analysis of activated neurons in the rat cortex. Error bars represent S.E.M. *p < 0.05 compared with sham control, two way analysis of variance on ranks with Dunn’s method. Scale bars: B, 200 μm

**3. Supplementary References**

There is no specific additional references in supplementary document.
