## Supplementary material for "Effect of sinusoidal electrical cortical stimulation on brain cells": highlights

- In vivo, Inhibitory neurons were mainly activated at 20 Hz and excitatory neurons at 40 and 60 Hz
- In silico, Sinusoidal EBS induced excitatory and inhibitory activation was widest at 20 Hz of EBS
- Sinusoidal EBS had a major effect on inhibitory neurons and related neural circuits
